# Intron-mediated enhancement is not limited to introns

**DOI:** 10.1101/269852

**Authors:** Jenna E Gallegos, Alan B Rose

## Abstract

Certain introns strongly increase mRNA accumulation by a poorly understood mechanism known as Intron-Mediated Enhancement (IME). Introns that boost expression by IME have no effect when located upstream of or more than ~1 Kb downstream from the start of transcription. The sequence TTNGATYTG, which is over-represented in promoter-proximal introns in *Arabidopsis thaliana*, can convert a non-stimulating intron into one that strongly increases mRNA accumulation. We tested the ability of an intron containing this motif to stimulate expression from different locations and found that it had the same positional requirements as naturally occurring IME introns. The motif also stimulated gene expression from within the 5’-UTR and coding sequences of an intronless construct. Furthermore, the 5’-UTR of another gene increased expression when inserted into an otherwise non-stimulating intron in coding sequences. These results demonstrate that splicing is not required for intron-mediated enhancement, and suggest that other sequences downstream of the transcription start site in addition to introns may stimulate expression by a similar mechanism.

## Introduction

Significant efforts have been invested in identifying the DNA sequences that control the expression of individual genes in eukaryotes. These studies have revealed many common kinds of regulatory elements that collectively constitute promoters in the broadest sense of the term. These include the sites surrounding and immediately upstream of the transcription start sites (TSSs) to which general transcription factors bind to form the pre-initiation complex, proximal binding sites (usually less than 1kb from the TSS) for regulatory transcription factors, and distal elements, such as enhancers, which can affect expression over great distances in either direction (reviewed in [1–5]).

In addition to these well-known regulators of transcription, other transcribed sequences can play an important role in controlling expression. 5’ and 3’ UTR’s have been shown to influence mRNA stability, export, and translation (reviewed in [6–8]), exons can contain transcription factor binding sites [9] or intragenic enhancers [10], and introns have been shown to affect gene expression by a number of known and unknown mechanisms.

Some introns contain enhancers [11,12], alternative transcription start sites [13], or transcription factor binding sites [14]. In addition, splicing can have a general positive effect on expression via coupling with other mRNA processing events such as capping and polyadenylation [15]. Deposition of the exon junction complex proteins also aids in mRNA export and translation [16,17], and splicing can influence transcription by affecting the phosphorylation state of RNA polymerase II [18].

Certain introns can increase gene expression by an additional, poorly understood mechanism referred to as intron mediated enhancement (IME) [19]. Several properties of IME indicate that these introns influence expression in a manner that is mechanistically distinct from enhancers or proximal promoter elements. The most thoroughly analyzed IME intron is the first intron from the Arabidopsis *UBQ10* gene. This intron increases mRNA accumulation only when located downstream of the TSS, and can stimulate expression from at least 550 nt from the TSS but not 1,100 nt or more[20,21]. Remarkably, deleting a 300 nt region of the proximal promoter that contains all known TSSs does not diminish the expression of constructs containing this intron [21]. Transcription in these promoter-deleted constructs initiates in normally untranscribed sequences the same distance upstream of the intron as when the promoter is intact. The observation that many introns stimulate gene expression only from within transcribed sequences near the 5’-end of a gene [20,22–24] is the basis for the IMEter algorithm described below.

The role of intron splicing in IME is a subject of some debate [19]. Splicing is clearly not sufficient for IME, because many efficiently spliced introns have no effect on expression [25]. Testing whether or not splicing is necessary for IME is complicated by the fact that disrupting splicing has many consequences. Constructs with an unspliceable intron produce mRNA that differs in size and structure from constructs containing a spliceable intron or an intronless control. Thus mRNA with a retained intron may differ in stability or translatability. The intron sequences retained in the mRNA can also cause frame shifts or contain premature start or stop codons, all of which might abolish translation of the reporter gene and lead to mRNA instability through nonsense-mediated mRNA decay. In cases where splicing was prevented but the reading frame was preserved by adjusting intron length and eliminating in-frame start and stop codons, expression levels were reduced but not eliminated [25–28]. The degree to which expression levels dropped varied greatly by species and the size, location, and original stimulating ability of the intron, precluding broad conclusions about the need for splicing in IME.

The differing ability of spliced introns to increase mRNA accumulation implies that some must contain stimulating sequences that others lack. These sequences have proven difficult to identify because they are redundant and dispersed throughout stimulating introns [26,29]. Progress was made using the IMEter algorithm, which generates a score that reflects the degree to which the oligomer composition of a given intron resembles that of promoter-proximal introns genome-wide. High IMEter scores have accurately predicted the stimulating ability of introns in Arabidopsis [29], soybeans [30], and other angiosperms [31]. The IMEter does not directly reveal stimulating sequences but can be used to identify sufficient numbers of potentially stimulating introns to allow computational searches for shared sequences.

One such motif, TTNGATYTG, was found to be over-represented in introns with high IMEter scores [29]. Rearranging nucleotides to create 6 or 11 copies of this motif converted a non-stimulating intron from the *COR15a* gene in Arabidopsis into one that boosts mRNA accumulation 14- or 24-fold, respectively [32]. Introns containing this motif behave similarly to the *UBQ10* intron in that they increase mRNA accumulation even in the absence of the proximal promoter [21], suggesting that the sequence TTNGATYTG is sufficient for IME.

The identification of the TTNGATYTG motif provided an opportunity to determine whether the mechanism of IME requires splicing, and is therefore specific to introns, or if it could act to increase expression from any location within the first few hundred nucleotides of transcribed sequences. We therefore tested the ability of this sequence to stimulate gene expression from within promoter-proximal exon sequences of an intronless construct. We found that six copies of the motif in exon sequences stimulate mRNA accumulation almost as much as does an intron containing six copies of the motif. This demonstrates that splicing is not required for IME, and that the sequences that increase expression by an IME mechanism can also function in exons near the start of a gene.

## Results

### The TTNGATYTG motif stimulates expression over a limited range

To first determine if the TTNGATYTG motif has positional requirements similar to the *UBQ10* and other natural introns, the ability of an intron engineered to contain 11 copies of the motif to stimulate expression was tested from five locations within a *TRP1:GUS* reporter construct. This intron, which was previously generated and designated *COR15a*11L [32], was created by rearranging sequences within a naturally non-stimulating intron from the *COR15a* gene [25,29]. The *COR15a*11L intron was inserted into *TRPI:GUS* fusion constructs at one of five locations: upstream of the transcription start site, within the 5’ UTR, at the 3’ end of sequences derived from *TRP1* exon 1, near the middle of the *GUS* gene, or at the 3’ end of the *GUS* gene (Figure 1A).

**Figure 1:**
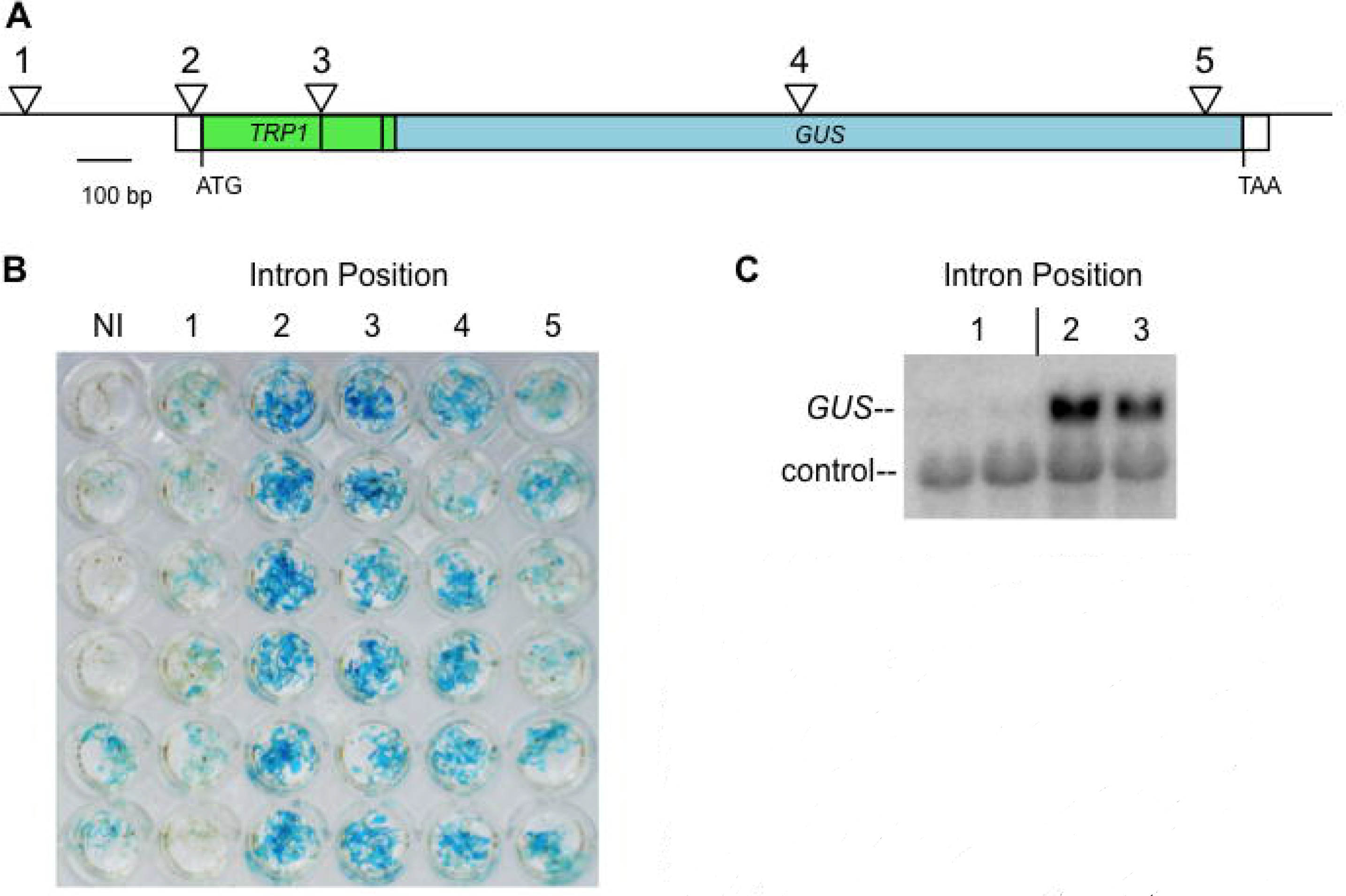
Activity of intrans containing the TTNGATYTG motif at different locations. **A)** The *TRP1:GUS* reporter gene, with triangles marking the sites where the *COR15a11L* intron was introduced. Shaded rectangles indicate protein coding sequences, open rectangles denote 5′ and 3′ untranslated regions. **B)** Histochemical staining for *GUS* activity in transgenic plants that contain the *TRP1:GUS* fusion with no intron (NI) or the *COR15a11L* intron at the posit on indicated at the top of each column. Each well contains five T_2_ seedlings from an independent transgenic line. **C)** RNA gel blot probed with *GUS* and a loading control (the endogenous *TRP1* gene). Each lane contains RNA from an independent single-copy homozygous line.

*TRPI:GUS* constructs containing the *COR15a*11L intron at the different locations were transformed into Arabidopsis and expression levels were compared. GUS activity was determined by histochemical staining of kanamycin-resistant T_2_ seedlings in lines of unknown copy number but whose segregation ratios indicated a single locus of insertion. The *COR15a*11L intron strongly stimulated expression from the 5’ end of the gene [32], but had no effect from upstream of the TSS and little effect from the 3’ end of the gene (Figure 1B).

To determine the level of expression responsible for the observed difference in GUS activity, RNA gel blots were performed on single-copy transgenic lines in which the intron was inserted near the TSS. The differences in GUS activity of these lines is reflected in the steady state *GUS* mRNA levels, indicating that the intron must be downstream of the TSS to increase mRNA accumulation (Figure 1C). The lack of stimulating effect of the *COR15a*11L intron from the 3’ end of the *TRP1:GUS* fusion or upstream of the TSS rules out the possibility that the TTNGATYTG motif acts as a conventional enhancer or transcription factor binding-site, and suggests that it increases mRNA accumulation by an IME mechanism.

### The TTNGATYTG motif stimulates expression from within exons in the absence of splicing

To determine if sequences involved in IME can stimulate expression in the absence of splicing, six copies of the sequence TTAGATCTG (the most active tested version of the TTNGATYTG motif [32]) were engineered into the first 450 nt of transcribed sequences of an intronless *TRP1:GUS* fusion (Figure 2). The *TRP1* sequences are described as exons 1, 2, or 3 based on their location in the endogenous *TRP1* gene. Five motifs were introduced into *TRP1* exon 1 sequences, two of which were in the 5’ UTR, and one copy was introduced into *TRP1* exon 2 sequences (Figure 3A). As a negative control, a second *TRP1:GUS* fusion was generated in which the AT dinucleotide at the center of each motif was changed to TA, making the motif TTAG<u>TA</u>CTG. This small inversion was previously shown to eliminate virtually all of the motif’s effect on mRNA accumulation from within an intron [32].

**Figure 2:**
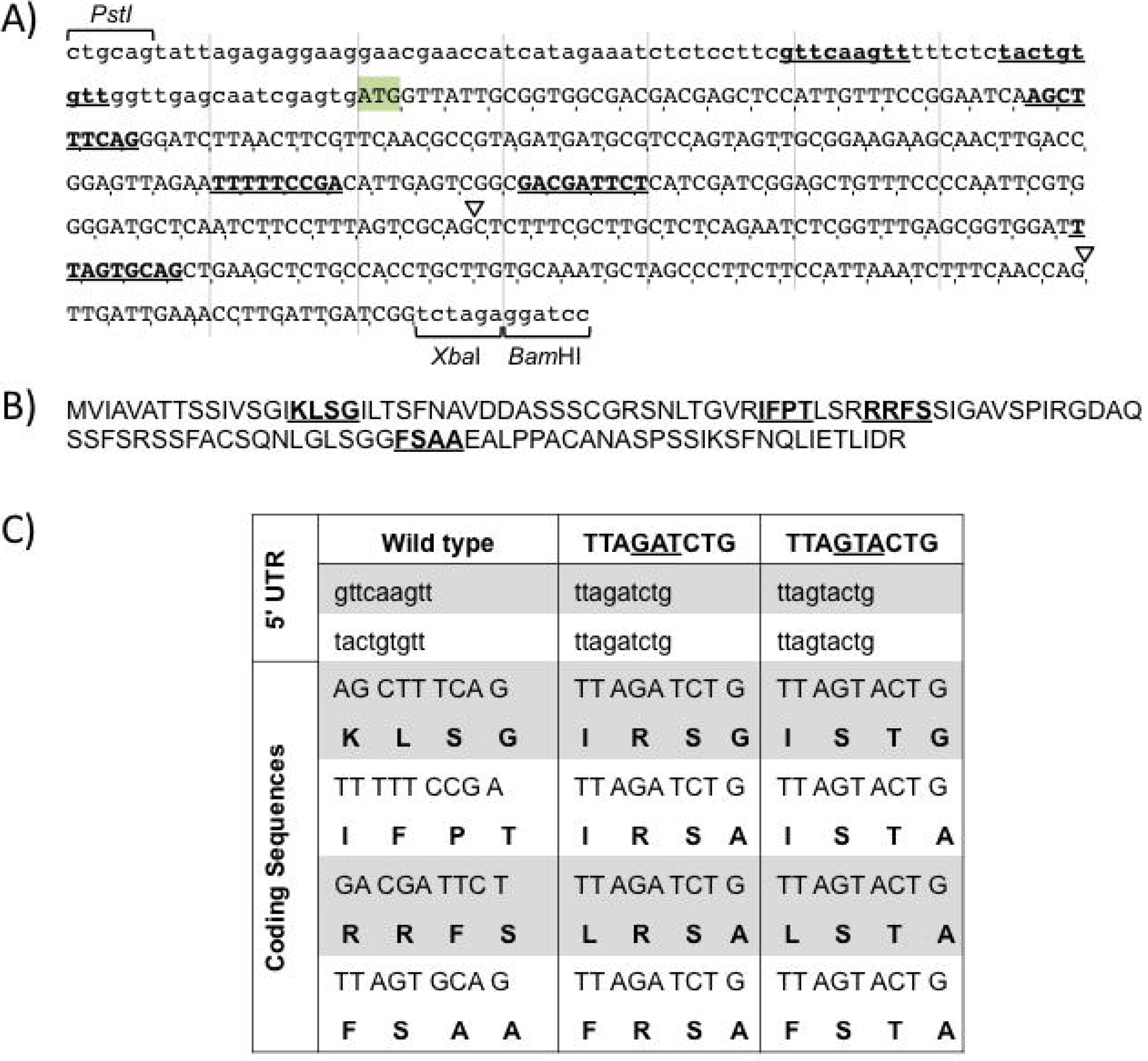
Details of changes to *TRPI:GUS* sequence to introduce motifs. **A)** Uppercase letters mark *TRPI* coding sequences with the start codon highlighted. Lowercase letters indicate the *TRPI* 5′ UTR and the sites used to fuse *TRP1* to *GUS*. The inverted triangles show the location of intrans in the endogenous *TRP1* gene. Sequences that are underlined and bold were changed to either TTAG<u>AT</u>CTG or TTAG<u>TA</u>CTG. **B)** The amino acids encoded by the *TRPI* sequence shown in A, with the regions affected by introducing the motifs underlined and in bold. **C)** Details of the nucleotides and amino acids changed.

Locations for introducing the motifs were selected to minimize changes to mRNA and protein structure (Figure 2). Existing sequences were searched for nine contiguous nucleotides composed of two As, one C, two Gs, and four Ts. Sequences that matched this criterion, or differed by no more than two nucleotides, were rearranged into the sequence TTAG<u>AT</u>CTG or TTAG<u>TA</u>CTG. In this way, the mRNAs from the tested constructs and controls would remain virtually unchanged in GC content and length. Locations were also selected to maximize the degree to which the changes in amino acids were conservative and consistent with the composition of chloroplast transit peptides. The first two exons of the *TRP1* gene encode a chloroplast transit peptide [33], which are rich in serines and threonines and usually devoid of negatively charged amino acids [34]. Chloroplast transit peptides are also poorly conserved and are cleaved off during chloroplast import [35]. Therefore, the changes made to introduce motifs were expected to have minimal effects on the activity of the mature GUS protein.

The stimulating ability of the motif in exonic locations was compared with that of the *COR15a6L* intron located between *TRP1* exon 1 and exon 2 sequences. The *COR15a6L* intron was previously generated by introducing six copies of the TTNGATYTG motif into the non-stimulating *COR15a* intron [32]. Expression was measured in single-copy transgenic Arabidopsis at both the mRNA and enzyme activity level and compared to intronless *TRP1:GUS* controls (Figure 3, Supplemental Tables 1 & 2).

**Figure 3:**
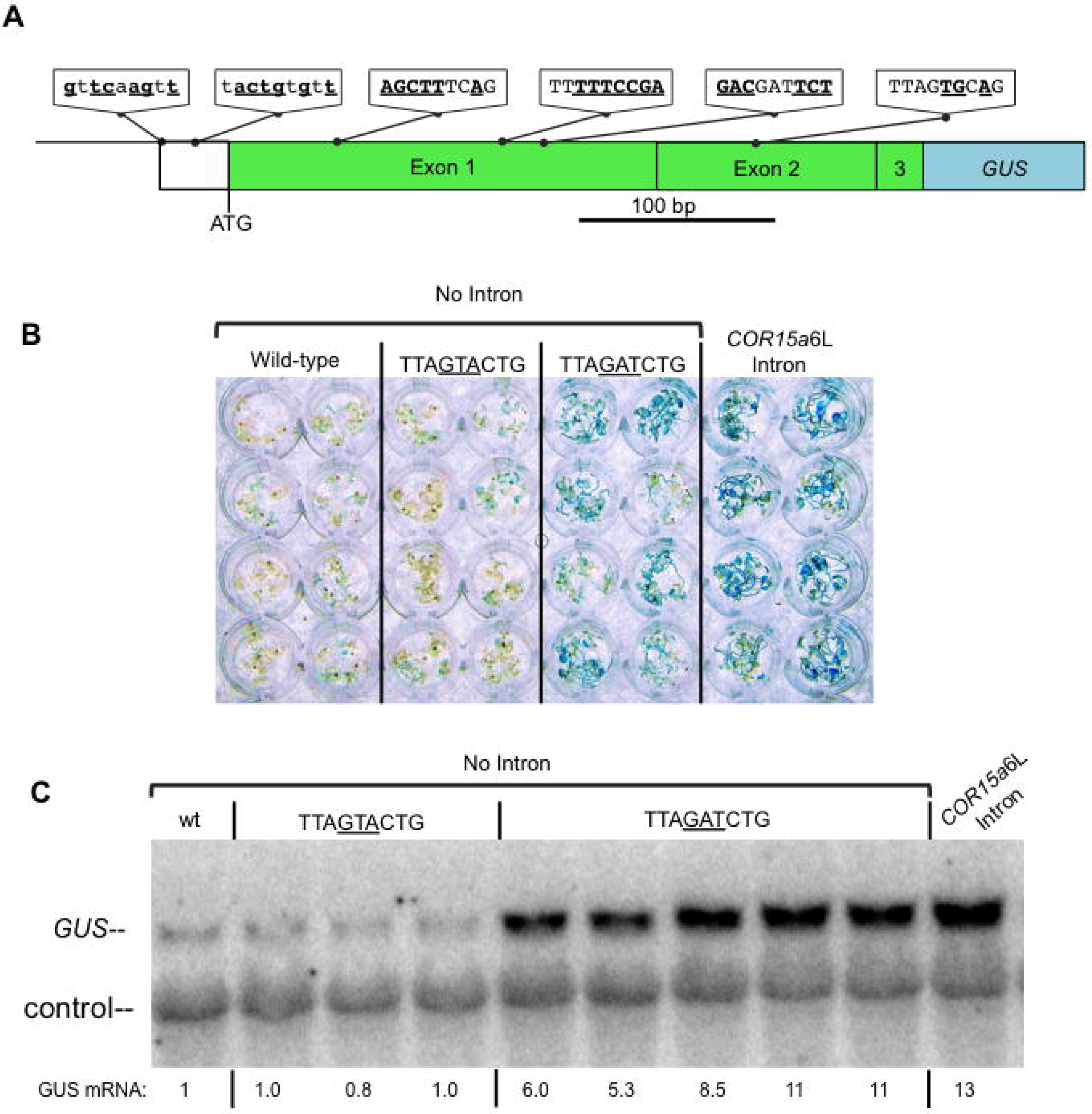
Testing the ability of the TTNGATYTG mot f to stimulate expression from within exons. **A)** Thenucleotides in the sequences at the indicated locations in a *TRP1:GUS* fusion that were changed to match the motif are bold and underlined. The designat ons of exons 1, 2, and 3 refer to the location of the same sequences in the endogenous *TRP1* gene. The *COR15a*6L intron in the control construct is located at thejunction between *TRP1* exon 1 and 2 sequences. No other constructs contain an intron. **B)** Histochemical staining for GUS activity in transgenic plants that contain the indicated *TRP1:GUS* fusions. Each well contains five T_2_ seedlings from an independent line of unknown copy number **C)** RNA gel blot probed with *GUS* and a loading control (the endogenous *TRP1* gene). Each adjacent lane with the same label represents an independent single-copy homozygous line.

The intronless *TRP1:GUS* fusion containing six copies of the TTAG<u>AT</u>CTG motif in exons showed substantially more histochemical staining for GUS activity than either the unmodified intronless control or the construct containing the mutated motif (Figure 3B, Supplemental Table 2). Single copy lines containing *TRP1:GUS* fusions with the TTAG<u>AT</u>CTG motif in exons accumulated on average 7.4 times more *TRP1:GUS* mRNA than did the unmodified intronless control (Figure 3C, Supplemental Table 1). This is slightly less than the 9.3 fold increase in expression caused by six copies of the same motif within the *COR15a*6L intron. In contrast, the fusion containing the mutated motif produced about the same amount of mRNA as the unmodified intronless control. Therefore, the increase in mRNA accumulation caused by the motif is similar in intronic and exonic locations, indicating that the TTAGATCTG motif can boost expression in the absence of splicing.

### 5’-UTR sequences can stimulate expression from within an intron

The ability of intron sequences to affect expression from the 5’-UTR, and the observation that average IMEter scores are nearly as high for 5’-UTRs as they are for promoter-proximal introns, supports the idea that sequences other than introns might boost expression by an IME mechanism [36]. To test this idea, we identified a 5’ UTR with a high IMEter score and examined the ability of this sequence to increase expression when inserted into a non-stimulating intron in a *TRP1:GUS* fusion.

The 5’-UTR chosen was a 74 nt fragment from the *At5g53000* gene, which encodes the Tap46 regulatory subunit of protein phosphatase 2A [37]. This fragment was used to replace an 80 nt internal portion of the *COR15a* intron, which was located at the 3’ end of *TRP1* endogenous first exon of a *TRP1:GUS* fusion.

The expression of each construct and an unmodified intronless control was measured as *TRP1:GUS* mRNA and GUS enzyme activity in single-copy transgenic Arabidopsis. The *At5g53000* 5’-UTR fragment stimulated mRNA accumulation more than 2-fold from within the previously non-stimulating *COR15a* intron (Figure 4, Supplemental Table 1). The effect of the 5’-UTR fragment on GUS enzyme activity (5.7-fold) was roughly twice that seen at the level of mRNA accumulation, as has been observed previously for introns in *TRP1:GUS* constructs (20, 25, 32). Therefore, the high IMEter score of this 5’-UTR fragment is biologically relevant. The ability of this non-intron sequence to stimulate expression from within an intron further supports the idea that sequences that increase expression by an IME mechanism may not be limited to introns.

**Figure 4:**
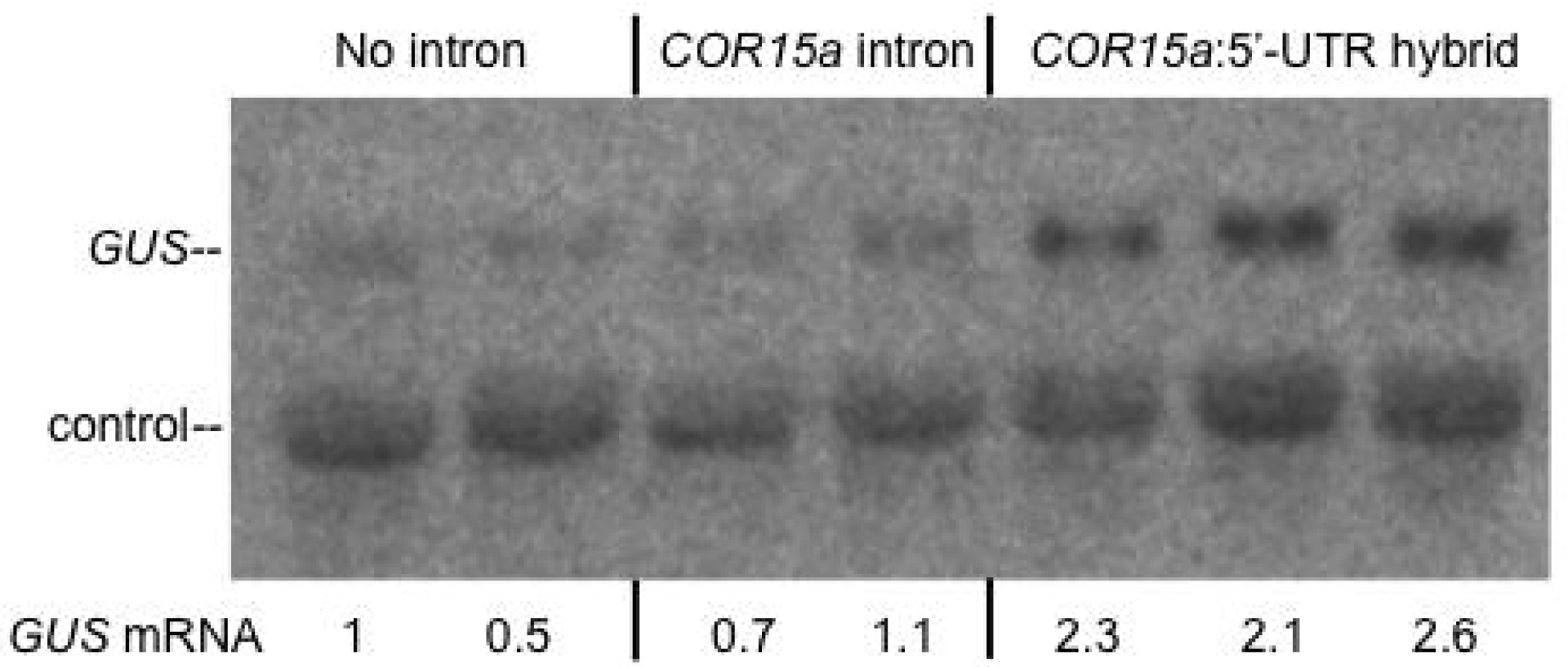
5’-UTR sequences can stimulate expression from within an intron. RNA gel blot of RNA from transgenic lines containing a *TRP1*:GUS fusion with no intron, the *COR15a* intron, or a hybrid intron in which a region of the *COR15a* intron was replaced with 5’-UTR sequences, probed with *GUS* and a loading control (the endogenous *TRP1* gene). Each ad acentlane with the same label represents an independent singlecopy homozygous line. Additional replicates are reported in Supplemental Table 1.

## Discussion

### Sequences associated with IME can contribute to mRNA accumulation in a position dependent and splicing independent manner

Multiple copies of the TTNGATYTG motif, which is overrepresented in introns with high IMEter scores, stimulate gene expression more than ten-fold from within a normally non-stimulating intron [32]. Here we showed that introns containing this motif strongly stimulated expression only from within transcribed sequences near the 5’ end of a gene. This locational specificity is a hallmark of introns that exhibit IME [20,21], further demonstrating that the TTNGATYTG motif is sufficient for IME. Because some introns that are known to stimulate expression in a position-dependent manner contain few matches to this motif, there must also be other sequences that act similarly.

Two lines of evidence suggest that IME is predominantly splicing-independent. First, the degree to which mRNA accumulation was increased by six copies of the TTNGATYTG motif was similar when the motifs were located within exons or in an intron. Second, IMEter scores, which strongly correlate with the ability of an intron to increase mRNA, are generally high in 5’-UTRs and to a lesser degree coding sequences near the start of a gene [36]. The ability of a high-scoring 5’UTR fragment from the *At5g53000* gene to stimulate mRNA accumulation from within the previously non-stimulating *COR15a* intron suggests that high IMEter scores of promoter-proximal non-intron sequences might also reflect their potential to increase mRNA accumulation by the same mechanism.

Even though the TTAGATCTG motifs increased mRNA accumulation from locations outside of introns, the effect of those sequences was somewhat higher when they were located within an intron. Thus, while the ability of intron sequences to stimulate gene expression is predominantly splicing independent, splicing may also contribute to an increase in mRNA accumulation. The effect of splicing on mRNA levels could be due to known interactions between the splicing and transcription machinery that increase transcription initiation or elongation, or synergies between splicing and other steps of mRNA processing. The overall level of expression is likely determined by multiple mechanisms, some of which are splicing-dependent and some of which are sequence-dependent.

The observations that intron sequences can stimulate gene expression from coding sequences outside of the context of an intron, that a 5’-UTR sequence with a high IMEter score can increase mRNA accumulation from within an intron, and that average IMEter scores are high genome-wide in 5’-UTRs, suggests that the mechanism through which expression is stimulated might include other sequences near the 5’ end of genes. The IMEter may thus be a useful tool for identifying potentially stimulating sequences in both exons and introns.

### Possible mechanisms

The properties of the described phenomenon are difficult to reconcile with known mechanisms of gene regulation. Many of the proposed mechanisms for IME such as gene looping or antisense suppression are predicted to require splicing [38,39], while the specific phenomenon reported here, which relates to high IMEter scores, appears to be predominantly splicing-independent.

It is unlikely that mRNA stability can account for the effect these sequences have on expression. Predictions of RNA structure reveal no obvious differences between the folding of *TRP1* sequences modified to contain the stimulating TTAG<u>AT</u>CTG motif compared with the non-stimulating TTAG<u>TA</u>CTG derivative. Additional lines of evidence further support a DNA-based mechanism for IME [21,40].

Several DNA-based mechanisms remain consistent with the results presented here and elsewhere, although some aspects remain puzzling for each. The ability of the TTNGATYTG motif to stimulate mRNA accumulation, while small changes to this sequence reduce or eliminate its effect on expression [32], suggests that there may be a protein such as a transcription factor that binds the motif in a sequence-specific manner. However, such a transcription factor would be unique for genes transcribed by RNA polymerase II in that it only functions when downstream of the transcription start site, and activates transcription several hundred nucleotides upstream of its binding site. The TTNGATYTG motif most closely resembles consensus sites for the GATA family of transcription factors [41,42], but GATA factor-binding sites do not meet the strict positional requirements characteristic of IME [43–45]. A second possible DNA-based model might include effects on local chromatin structure that favor transcript initiation, but this would not explain why these sequences must be downstream of the TSS to stimulate expression. A third possibility is that IME sequences influence the transcription machinery during elongation, elevating mRNA production by increasing processivity or the rate of transcription. This mechanism would not account for the ability of these sequences to boost expression in the absence of prior promoter activity [21].

Whatever the mechanism, it is clearly distinct from known effects of splicing, conventional transcription factor binding sites, and enhancer elements.

### Implications in gene evolution and codon usage bias

Introns with high IMEter scores are associated with strongly expressed constitutive genes. It is possible that housekeeping genes have evolved sequences throughout their 5’ ends that maximize ubiquitous expression. These sequences may be more commonly identified in introns due to the relative ease of generating and studying cDNA. Similar experiments exploring the effects of coding sequences and 5’-UTRs on expression are difficult to perform without introducing multiple confounding variables. Introns may also be more likely to contain stimulating sequences because they are under fewer evolutionary constraints than 5’-UTRs and coding sequences. However, the degenerate genetic code does allow for some flexibility in coding sequences. Not all codons are used with the same frequency, and this codon-usage bias can have dramatic effects on gene expression by diverse mechanisms (reviewed in[46]).

N-terminal codon selection is thought to be especially important in determining expression levels. Effects on RNA secondary structure are the largest contributing factor, but still only explain about half of the variation observed [47,48]. In yeast, synonymous mutations at the 5’ ends of genes have been shown to impact nucleosome positioning [49]. Synonymous substitutions also appear to occur less frequently at the 5’ end of genes in mammalian populations (as determined by comparing evolution of the *BRCA-1* gene in humans and dogs)[50]. In addition to codon usage, nucleotide frequency distributions also differ along the lengths of genes, suggesting that promoter proximal sequences may have evolved in response to pressures such as maximizing gene expression [51–53]. Further, optimizing expression by varying codon usage is more effective when adjacent codon pairs, rather than individual codons, are considered [54–56]. It is possible that some of the observed variation in expression associated with codon-usage bias is due to the inadvertent creation or destruction of stimulating sequences in coding regions.

### Conclusion

What had been previously characterized as intron-mediated enhancement may not be limited to introns, and intron-mediated enhancement appears to be at least partially splicing independent. The ability of transcribed sequences near the start of genes to affect mRNA accumulation extends beyond introns and may include 5’-UTRs or coding sequences. These transcribed expression-stimulating sequences can be a useful addition to the promoters and enhancers used to regulate gene expression levels in transgenic or synthetic constructs.

## Materials and Methods

### Cloning of reporter gene fusions

The starting intronless *TRP1:GUS* template for all constructs included a 2.4 Kb *TRP1* promoter fragment that extends from the middle of the upstream gene (At5g17980) through the first 8 amino acids of the third exon of *TRP1* fused to the *E. coli uidA* (*GUS*) gene in the binary vector pEND4K [57]. To test the ability of the previously generated *COR15a11L* intron to stimulate expression from four additional locations (Figure 1), the intron, which is flanked by *PstI* sites [19], was cloned into previously generated *TRP1:GUS* constructs with *PstI* sites either 1136 or 1875 nucleotides downstream of the major transcription start site [20], or 21 or 324 nucleotides upstream of the *TRP1* start codon [21].Transgenic plants in which the *COR15a11L* intron is located between the endogenous first and second exons of *TRP1:GUS* were previously described [32]. Other introns flanked by *PstI* sites are efficiently spliced at these locations [20,21,32].

To introduce the TTAG<u>AT</u>CTG motif and TTAG<u>TA</u>CTG control motif into exons (Figure 2), *TRP1* sequences containing the described changes were synthesized by Biomatik (Wilmington, Deleware) and confirmed by sequencing. These fragments were used to replace analogous sequences between a *PstI* site engineered into the *TRP1* 5’ UTR 87nt upstream of the start codon [21] and a *BamHI* site in the polylinker region connecting the *TRPI* and *GUS* coding sequences.

A 74 nucleotide *BamHI* fragment from the 3’ end of the 5’ UTR of *A5tg53000* was used to replace an 80 nt *Bam*HI to *Bcl*I fragment of a modified version of the *COR15a* intron [29]. The inserted fragment from the *At5g53000* 5’ UTR differs from the analogous endogenous fragment by three nucleotides. Changes were generated to introduce restriction sites via PCR mutagenesis:

Original: t<u>**t**</u>caaaagacgatcctcttctcgaagaaactcgattcttgtggattcgatttcattaaggaattttgaattg<u>**tt**</u>
Inserted: t<u>**c**</u>caaaagacgatcctcttctcgaagaaactcgattcttgtggattcgatttcattaaggaattttgaattg<u>**ga**</u>

The resulting hybrid intron was then cloned as a *PstI* fragment into a previously generated *TRP1:GUS* construct with a *PstI* site located between the endogenous first and second exon of *TRP1* [20].

All fusions were then transformed into *Agrobacterium tumefaciens by* electroporation and introduced into *Arabidopsis thaliana* ecotype Columbia (Col) by floral dip as described [28].

### Qualitative GUS expression assays

To compare the effect of the *COR15a11L* intron on *TRP1:GUS* expression from various locations (Figure 1B), five T_2_ seedlings from each of multiple lines whose segregation ratios indicated a single locus of transgene insertion were histochemically stained for *GUS* activity. To compare the enzyme activity of constructs containing the TTNGATYTG motif from within exons and introns (Figure 3B), T_2_ seedlings from 12 lines of unknown copy number for each construct and controls (intronless control: pAR281 [28], *TRP1:GUS* with *COR15a6L* intron: pAH3 [32]) were histochemically stained for GUS activity. In both cases, the plate of seedlings in buffer (10mM EDTA, 100mM NaPO4 pH 7.0, 0.1%Triton X-100) containing 0.5 mg/mL 5-bromo-4-chloro-3-indolyl ß-D-glucuronic acid (Calbiochem, La Jolla, CA, USA) was incubated at 37° for one hour. The seedlings were washed in water and soaked in ethanol to remove chlorophyll.

### Quantitative comparisons of enzyme activity and mRNA levels

Single-copy transgenic lines were identified for several key constructs, and mRNA levels on RNA gel blots and GUS activity in leaf extracts were measured as previously described [20]. In short, seeds from several dozen lines were screened for a 3:1 segregation ratio (kanamycin resistant: sensitive), and gel blots of DNA digested with restriction enzymes were probed with the *GUS* gene to determine transgene copy number. Single copy, homozygous lines were propagated to the T_3_, T_4_, or T_5_ generation. RNA was extracted from 3-week-old seedlings, grown under constant light in Professional Growing Mix (Sun Gro Horticulture, Agawam, MA) at a density of 500 plants per 170 cm^2^ pot, using the Qiagen RNeasy kit. RNA gel blots were hybridized with a ^32^P-labeled *GUS* probe, and *GUS* mRNA levels in PhosphoImager scans were measured as pixels above background using Image Quant Software as described previously [29]. Quantitative measurements of GUS enzyme activity in leaf extracts were performed as described [20]. All mRNA and enzyme activity levels were normalized for total mRNA/protein and compared to the intronless control pAR281 [28].

Statistical differences in gene expression between constructs were analyzed by comparing Log mRNA levels using a mixed model that accounted for blot-to-blot differences, and adjusted for random effects per line and date of mRNA extraction for biological replicates. Residual normality was analyzed using a Wilk Shapiro test and homoscedasticity using a Levene ANOVA. Among the levels of categorical predictors, post hoc comparisons were based on least squares means using a protected least significant difference.

